# Recombinase-Independent AAV for Anterograde Transsynaptic Tracing

**DOI:** 10.1101/2023.06.20.545789

**Authors:** Islam Faress, Valentina Khalil, Haruka Yamamoto, Szilard Sajgo, Keisuke Yonehara, Sadegh Nabavi

## Abstract

Viral transsynaptic labeling has become indispensable for investigating the functional connectivity of neural circuits. Adeno-associated virus serotype 1 (AAV1) allows for anterograde transneuronal labeling and manipulation of postsynaptic neurons. However, it is limited to delivering an AAV1 expressing a recombinase which relies on using transgenic animals or genetic access to postsynaptic neurons. We reasoned that a strong expression level could overcome this limitation. To this end, we used a self-complementary AAV1 (scAAV1) under a strong promoter (CAG). We demonstrated the anterograde transneuronal efficiency of scAAV1 by delivering a fluorescent marker in mouse retina-superior colliculus and thalamic-amygdala pathways in a recombinase-independent manner. In addition to investigating neuronal connectivity, scAAV1.CAG may be suitable for functional manipulation and imaging.

## Introduction

Significant progress has been made in the fields of neurophysiology and neuroanatomy since the discovery of viral approaches to map out brain connectivity. The viral approach is a powerful tool to label interconnected brain regions, allowing cell-type-specific identification and manipulation. Retrograde mapping of the presynaptic inputs to a specific brain region is achievable via different viral tools that vary in their degree of specificity (axonal vs. transsynaptic transport) and efficiency (Luo et al., 2018; Tervo et., 2016). Anterograde tracing has remained largely limited to labeling axonal input to the downstream postsynaptic region. However, the recent discovery of the adeno-associated virus serotype 1 (AAV1) property to label postsynaptic neurons transsynaptically advanced the field. AAV1 expressing a recombinase has been used in various studies to label and manipulate postsynaptic neurons. The major limitation is that a recombinase-dependent transgene should be expressed in downstream targets, which is doable only via transgenic animals or viral expression postsynaptically, which requires prior knowledge of the target (Luo et al., 2018; Zingg et al., 2017; Zingg et al., 2020). This study aims to optimize a method allowing recombinase-independent anterograde transsynaptic labeling of postsynaptic neurons.

## Results

We optimized the expression parameters to achieve stronger expression of an AAV1 vector that carries a green fluorescent protein (GFP). The vector is self-complementary, I.e., carrying double-stranded DNA and under short-CAG as a promoter at a high titer (>1.0 × 10E13 vg/ml) (McCarty et al., 2001). To test the efficiency of the vector, we conducted two separate experiments in well-characterized long-range monosynaptic and non-reciprocal visual and auditory pathways. In the first experiment, we injected the vector into the mouse retina and imaged the downstream target in the superior colliculus (SC). In the second experiment, the virus was injected into the medial geniculate nucleus (MGn), and we imaged the basolateral amygdala (BLA). We also evaluated whether an AAV1 vector with an alternative (and possibly weaker) promoter, human Synapsin (hSyn), would yield a comparable level of transsynaptic anterograde labeling.

We delivered the scAAV1 GFP under either short.CAG or hSyn to the retina, and 3 weeks later, we harvested the brains and retinae to examine the GFP labeling postsynaptically (Fig.1 A-H). The two vectors were delivered separately at an equal titer and volume. We observed that scAAV1.short.CAG.GFP resulted in a significantly higher number of transsynaptically labeled GFP neurons in the SC, in comparison to scAAV1.hSyn.GFP (Fig.1 C-E). In the mice injected with scAAV1.short.CAG.GFP in the retina, we observed that the contralateral SC had robust neuronal GFP labeling, while the ipsilateral had very sparse labeling, which is consistent with basic retinal-collicular anatomy (Fig.1 F,G). Unlike the efficient transsynaptic anterograde labeling we observed in the SC from the retina, we did not observe any somatic labeling in the lateral geniculate nucleus (LGN). This lower labeling efficiency in the LGN compared to the SC is consistent with the original study where AAV1 has been reported to show anterograde transsynaptic labeling properties (Zingg et al., 2017). Next, we tested the intracerebral efficiency by injecting the scAAV1.short.CAG.GFP in the MGn.This resulted in robust labeling only in the ipsilateral BLA, and no labeling in the contralateral BLA, which is consistent with the known neuroanatomy (Fig.1 I-L) (Barsy et al., 2020; Hintiryan et al., 2021).

**Figure 1.**
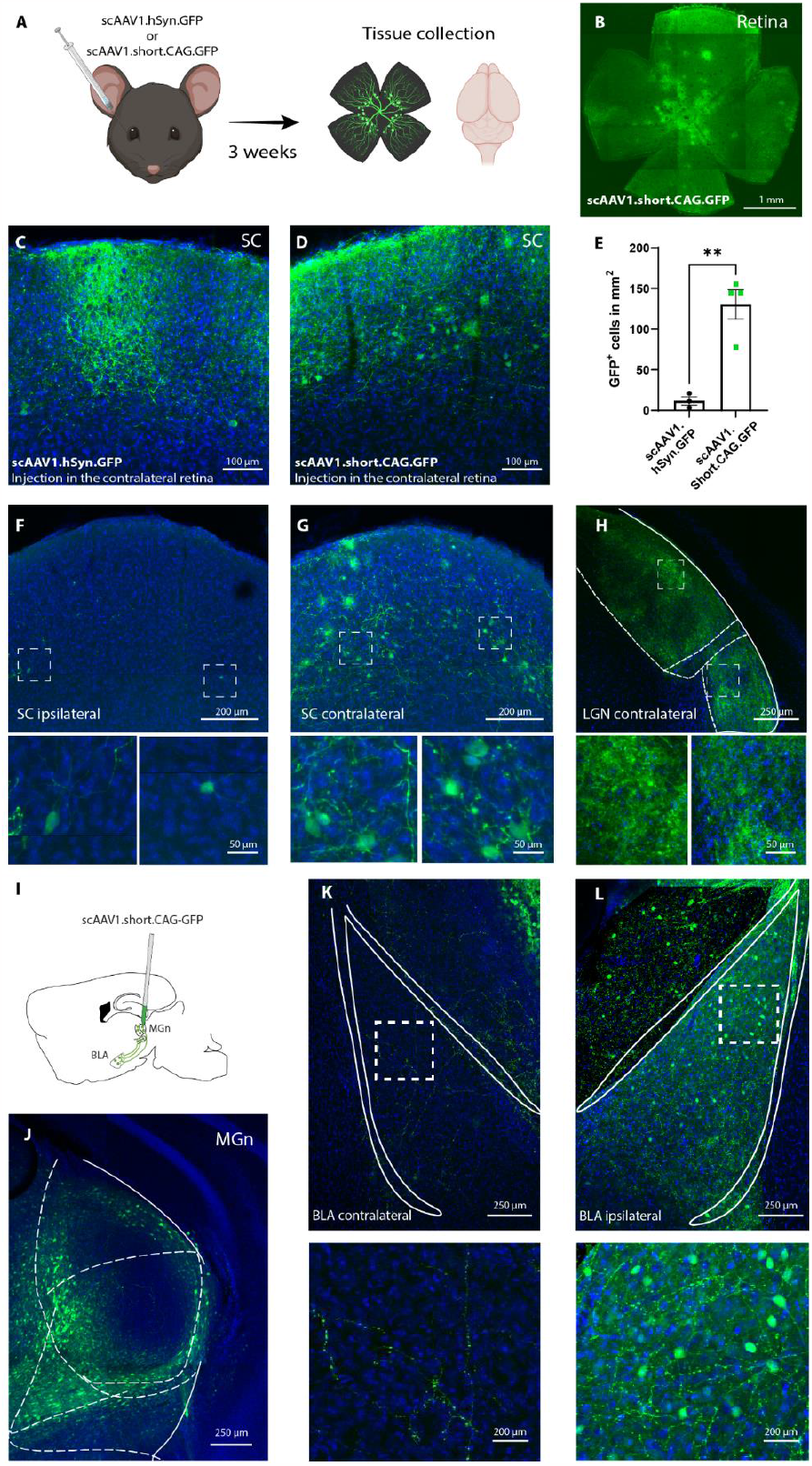
**A)** Diagram showing the experimental timeline for the retinal injection of either scAAV1.hSyn.GFP (n=3) or scAAVI .short.CAG.GFP (n=4). **B**) Image of a retina injected with scAAV1.short.CAG.GFP. **C, D**) Representative image showing GFP-labeled neurons in the SC after scAAV1.hSyn.GFP (**C**) and scAAV.1 short.CAG.GFP (**D**) retinal injection. **E**)The number of GFP-positive neurons in the SC is significantly higher in mice with scAAVI .short.CAG.GFP retinal injection (n=4) compared to mice with scAAVI .hSyn.GFP retinal injection (n=3; Unpareid t-test, p=0.0027). **F, G, H**) Representative images from the ipsilateral SC (**F**), contralateral SC (**G**), and LGN (**H**).The insets below are high-magnification images of the boxed regions. **I**) Diagram showing the injection of AAV.1 short.CAG.GFP in the MGn and the anterograde transsynaptically labeled neurons in the BLA (n-4). **J**) Representative image of the injection site in the MGn. **K**,**L**) Representative images showing that the ipsilateral BLA has GFP-positive neurons (**L**) after AAV1.short.CAG.GFP in the MGn, while the contralateral BLA (**K**) does not. The boxes below are high magnification of the respective regions. Results are reported as mean ± S.E.M. **, p<0.01.

## Discussion

Here we investigated the possibility of achieving anterograde transsynaptic labeling in a recombinase-independent manner using an AAV1 vector. We demonstrated that scAAV1.short.CAG.GFP could transneuronally label downstream postsynaptic targets with a single virus injection in wild-type mice. After injecting the vector into the retina and the MGn, we observed reliable labeling in the SC and the BLA, respectively. Single-stranded AAV1 CAG GFP was reported to lack transneuronal anterograde labeling capacity, which might be due to the low expression level (Zingg et al., 2017). Consistent with this hypothesis, we observed that scAAv1.short.CAG.GFP yields more robust transsynaptic labeling in comparison to a possibly weaker promoter like scAAV1.hSyn.GFP. Our results indicate that the transsynaptic transfer of transgenes that express optogenetic/chemogenetic actuators might be achievable by using the AAV1 expressing the transgene of interest. These viral tools will expand the potential utility of viral approaches for circuit interrogation on the pathway and cell-type specific manner. Future studies will be needed to optimize AAV-mediated transsynaptic tracing to allow for strict anterograde labeling and, thereby, make it applicable in reciprocally connected brain regions. Similarly, optimizations that allow for using AAV1 at a lower titer while maintaining the anterograde transsynaptic property might reduce the toxicity at the injection site.

## Materials and methods

### Subjects

All the procedures were performed on C57BL/6JRJ wildtype (Janvier, France). Mice were naïve and acclimated to the vivarium for at least a week before the beginning of the experiment. The mice were six to eight weeks old at the beginning of the experimental procedures. Animals were group-housed (3-4 per cage) with enriched conditions in a 12h light/dark cycle (the light switches on at 6 A.M.) with constant levels of humidity and temperature (22±1). Food and water were provided ad libitum. Experiments were conducted at Aarhus University in the Biomedicine department, Ole Worms Allé 8, Aarhus 8000. All the experimental procedures were conducted according to the Danish Animal Experiment Inspectorate.

### Retinal and brain injections

For intravitreal viral injections, mice were anesthetized with an intraperitoneal injection of fentanyl (0.05 mg/kg body weight; Actavis), midazolam (5.0 mg/kg body weight; Dormicum, Roche), and medetomidine (0.5 mg/kg body weight; Domitor, Orion) mixture dissolved in saline. We made a small hole at the border between the sclera and the cornea with a 30-gauge needle. 1.5 μl of the AAV vector was injected into the vitreous of the eye using a blunt-end microsyringe (Ito Corporation, #MS*E05). Mice were returned to their home cage after anesthesia was reversed by an intraperitoneal injection of flumazenil (0.5 mg/kg body weight; Anexate, Roche) and atipamezole (2.5 mg/kg body weight; Antisedan, Orion Pharma) mixture dissolved in saline. AAV delivery to MGn (0.3 μl), GFP immunohistochemical amplification, and quantification were done as described in (Khalil et al., 2023).

### Confocal imaging

Imaging was performed using the ZEN software (ZEN 2.5, blue edition) and a ZEISS confocal microscope (LSM 780/Axio Observer.Z1m). The images were taken as z-stack images (z-step size = 6 μm) in the respective areas using a 10× objective (Apochromat 10×/0.45 M27). Alexa 488 was excited at 488 nm and detected through a bandpass filter of 491–535 nm, while the DAPI was stimulated at 405 nm and detected through a 410–483 nm bandpass filter.

### Statistics

Statistical analyses were performed by using GraphPad Prism 9. All the data are represented as mean ± SEM, and they were tested for normality using Shapiro-Wilk. If the data represented a normal distribution, a parametric test was used. The statistical methods and the corresponding p-values are reported in the figure legend.

## Contributions

IF and SN conceived and designed the study. IF wrote the original draft with inputs from VK, HY, KY, and SN. IF, VK, KY, and SN designed the experiments. IF, VK, HY, and SS performed the experiments. VK made the figure.

## Acknowledgments

We thank members of the Nabavi laboratory for their suggestions. We thank Jean-Charles Paterna and Viral Vector Facility (VVF) of the Neuroscience Center Zurich (ZNZ) for their support. This study was supported by ERC starting grant (22736), by Independent Research (DFF), Novo Nordisk Foundation (NNF16OC0023368), and AUFF NOVA grants to SN. Additionally, SN was supported by the Danish Research Institute of Translational Neuroscience (19958), and by PROMEMO (Center of Excellence for Proteins in Memory funded by the Danish National Research Foundation) (DNRF133). HY was supported by The Lundbeck Foundation LF Postdocs (R347-2020-2299), Daiichi Sankyo Foundation of Life Science, and The Uehara Memorial Foundation research fellowship. Figures were Created with BioRender.com.

